# Reprogramming of Human Cells to Pluripotency Induces CENP-A Chromatin Depletion

**DOI:** 10.1101/2020.02.21.960252

**Authors:** Inês Milagre, Carolina Pereira, Raquel Oliveira, Lars E.T. Jansen

## Abstract

Pluripotent stem cells (PSCs) are central to development as they are the precursors of all cell types in the embryo. Therefore, maintaining a stable karyotype is essential, both for their physiological role as well as for use in regenerative medicine. In culture, an estimated 10-30% of PSC lines present karyotypic abnormalities, but the underlying causes remain unknown. To gain insight into the mitotic capacity of human embryonic stem cells and induced pluripotent stem cells, we explore the structure of the centromere and kinetochore. Centromere function depends on CENP-A nucleosome-defined chromatin. We show that while PSCs maintain abundant pools of CENP-A, CENP-C and CENP-T, these essential centromere components are strongly reduced at stem cell centromeres. Outer kinetochore recruitment is also impaired to a lesser extent, indicating an overall weaker kinetochore. This impairment is specific for the kinetochore forming centromere complex while the inner centromere protein Aurora B remains unaffected. We further show that, similar to differentiated human cells, CENP-A chromatin assembly in PSCs requires transition into G1 phase. Finally, reprogramming experiments indicate that reduction of centromeric CENP-A levels is an early event during dedifferentiation, coinciding with global chromatin remodelling. Our characterisation of centromeres in human stem cells drives new hypotheses including a possible link between impaired centromere function and stem cell aneuploidies.

## Introduction

Embryonic stem cells (ESCs) are derived from the inner cell mass and can give rise to all cell types in the embryo (Thomson et al., 1998). The maintenance of genome structure and ploidy is key to their ability to generate viable daughter cells and maintain their differentiation capacity. Despite their extensive proliferative potential, the mechanics of cell division in these cells are still under explored. One key component for faithful mitosis is the centromere, a specialized chromosomal locus that acts as a chromatin-based platform for the assembly of the kinetochore, composed of microtubule associated-proteins that drive chromosome segregation (Cheeseman and Desai, 2008). How centromere structure is maintained and how it is regulated in stem cells is still unknown.

Pluripotent stem cells can be of embryonic origin, however they can also be generated in culture using ectopic expression of only four transcription factors (Takahashi and Yamanaka, 2006) leading to the formation of induced pluripotent stem cells (iPSCs). These share various characteristics with ESCs, such as a truncated cell-cycle (Ghule et al., 2011), comparable cell morphology, self-renewal capacities, expression of pluripotency associated markers and the ability to differentiate into derivatives of all three primary germ layers (Takahashi and Yamanaka, 2006). The generation of iPSCs offers key tissue engineering opportunities and clinical applications. Additionally, they also represent a helpful tool in culture to understand how the stem cell state impacts on basic cell biology such as the mechanics of cell division and the fidelity of chromosome segregation.

Induction of pluripotency in differentiated cells requires the repression of somatic genes and activation of self-renewal and pluripotency associated genes. We and others have shown that reprogramming requires striking remodelling of chromatin modifications, such as global and targeted DNA demethylation at key regulatory regions (Lee et al., 2014; Milagre et al., 2017), including pluripotency related enhancers, super-enhancers (Milagre et al., 2017) and histone marks (Nashun et al., 2015). Specific histone marks, such as H3K4me2 and H3K9me3 are considered barriers to reprogramming as failure to remove or re-distribute these marks results in the inability of cells to reach pluripotency (Nashun et al., 2015). The profound remodelling of chromatin structure is what allows cells to transition from a somatic cell identity to a stable pluripotent cell identity, while maintaining the same genomic information. It is not clear how this genome-wide remodelling of the chromatin impacts on the structure and stability of the epigenetically defined centromere.

Both human ESCs and iPSCs appear to have an elevated level of genomic instability, at least in culture. Two reports have analysed hundreds of ESC and iPSC lines used in different laboratories worldwide and assessed that at around 10% to as much as 34% of all cell lines have abnormal karyotypes (International Stem Cell Initiative et al., 2011; Taapken et al., 2011). ESCs have a unique abbreviated cell cycle with a shortened G1 phase (Becker et al., 2006), and the rapid proliferation of these cells has been proposed both as a possible cause, but also as a consequence of these genomic abnormalities (Weissbein et al., 2014). Further, it has been shown that karyotypically abnormal pluripotent stem cells (PSCs) present defects in the capacity to differentiate into all cell types of the organism and display higher neoplastic capacity, thus hindering their potential application (Zhang et al., 2016). However, why these cells are prone to karyotypic defects is unclear.

Here we explore the structure of the centromere in both embryo-derived stem cells as well as induced pluripotent stem cells with the aim to understand the basis of mitotic fidelity and possible causes of aneuploidy. Central to the structure, function and maintenance of the centromere is an unusual chromatin domain defined by nucleosomes containing the histone H3 variant CENP-A (Black et al., 2010; McKinley and Cheeseman, 2016). Centromere specification is largely uncoupled from DNA cis elements (Marshall et al., 2008; Murillo-Pineda and Jansen, 2020) and maintenance depends primarily on a self-propagating CENP-A feedback mechanism (Hori et al., 2013; Mendiburo et al., 2011). We have previously shown in somatic cells that CENP-A is stably associated with chromatin throughout the cell cycle, consistent with a role in epigenetically maintaining centromere position (Bodor et al., 2013; Falk et al., 2015). CENP-A chromatin in turn recruits the constitutive centromere-associated network (CCAN) (Foltz et al., 2006; Okada et al., 2006). The key components of this network are CENP-C and CENP-T that make direct contacts to the microtubule binding kinetochore in mitosis (Gascoigne et al., 2011; Hori et al., 2008). CENP-A chromatin propagation is cell cycle regulated and restricted to G1 phase, through inactivation of the cyclin-dependent kinases (Cdk1 and Cdk2) (Silva et al., 2012; Stankovic et al., 2017). Nascent CENP-A is guided to the centromere by the HJURP chaperone in a manner dependent on the Mis18 complex (Barnhart et al., 2011; Dunleavy et al., 2009; Foltz et al., 2009), both of which are under strict cell cycle control (McKinley and Cheeseman, 2014; Stankovic et al., 2017).

Although the mechanisms of centromere assembly and the cell cycle control thereof are well established in somatic cells, virtually nothing is known about centromere regulation in PSCs. Here we define the composition and size of the human centromere in both ESCs as well as iPSCs and find that stem cells maintain a reduced centromeric chromatin size, impacting the key centromere proteins CENP-A, CENP-C and CENP-T, despite ample pools of cellular protein. This reduction in centromere size is recapitulated by induction of the stem cell state and coincides with early reprogramming.

## Results

### Pluripotent stem cells have a weaker centromere than differentiated cells

To characterize the mitotic performance of human embryonic stem cells (ESCs) we cultured the established embryonic stem cell line H9 (hESCs, henceforth) and determined the fidelity of chromosome segregation. To this end, we fixed and scored mitotic cells for chromosome segregation errors. We compared segregation rates to human RPE-1 cells (RPE, henceforth) as a representative immortalized somatic epithelial cell line. In agreement with previous reports (International Stem Cell Initiative et al., 2011; Taapken et al., 2011), we find that cultured human ESCs have a twofold elevation in total chromosome missegregation events (Figure S1A).

To characterize centromere size and function in ESCs we compared centromere protein levels by immunofluorescence in hESCs cells and RPE cells in which we have previously characterised centromeres in detail (Bodor et al., 2014). Furthermore, we reprogrammed human primary fibroblast derived from adult skin into induced pluripotent stem cells (iPSCs) by Sendai virus-mediated transduction of the Yamanaka reprogramming factors [Oct4, Sox2, Klf4 and c-Myc (Takahashi and Yamanaka, 2006)]. We reprogrammed fibroblasts from two different human donors to iPSCs, which express Sox2 and Nanog, to levels comparable to hESCs (Figure S1B). CENP-A containing nucleosomes form the chromatin platform upon which the centromere complex and kinetochore is build. Despite the essential nature of ESCs to life and development, we find centromeric chromatin to be greatly reduced in CENP-A nucleosomes numbers, at ∼40% of the levels observed in RPE cells (Figure 1A, B). Next, we determined whether reduced centromeric chromatin size is unique to hESCs or whether this is a general phenomenon across stem cells. In agreement with the data derived from embryonic stem cells, iPSCs also show a dramatic decline of CENP-A levels at the centromere, to as little as 25% of RPE levels and 29 to 42% of the levels observed in the donor fibroblasts (donor#2 and donor#1, respectively) from which the iPSCs were reprogrammed (Figure 1A, B, S1C). This latter result demonstrates that reduced centromeric CENP-A is directly linked to the epigenetically determined stem cell state as the iPSCs are genetically identical to their cognate donor fibroblasts. We confirmed these results by cell fractionation experiments. We observed that hESC have reduced levels of CENP-A in the chromatin bound fraction with a comparative increase in the soluble fraction, when compared to RPE cells (Figure S1C). Consistently, the ratio of chromatin bound to soluble pool of CENP-A, is decreased in hESC (Figure S1D, E).

**Figure 1.**
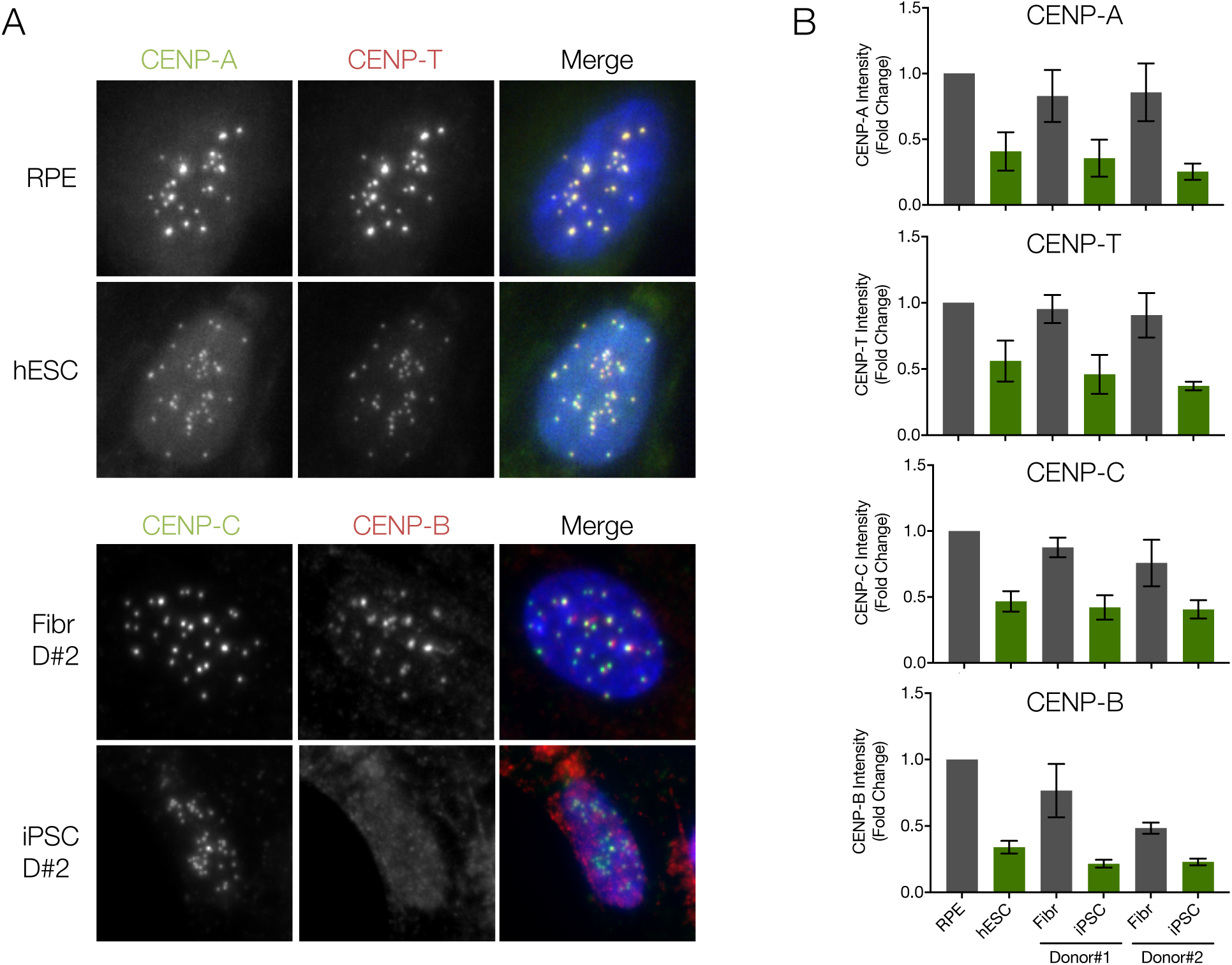
Pluripotent stem cells have a weaker centromere than differentiated cells. **A)** Differentiated (Retinal Pigment Epithelium – RPE and fibroblasts from two independent donors – Fibr D#1 and Fibr D#2) and pluripotent stem cells (human Embryonic Stem Cell line H9 – hESC or iPSC lines reprogrammed from fibroblasts from Donor #1 and Donor #2 – iPSC D#1 or iPSC D#2) were fixed and stained for CENP-A, CENP-T, CENP-C or CENP-B and counterstained with DAPI (blue). Representative immunofluorescence images from RPE and human embryonic stem cells (hESCs) are shown for CENP-A and CENP-T and representative images from Fibroblasts and iPSC from donor #2 are shown for CENP-B and CENP-C. **B)** Quantification of centromere intensities as shown in A) for all cell types. Average centromere intensities were determined using automatic centromere recognition and quantification (CRaQ; see methods). The average and standard error of the mean of three replicate experiments are shown. Centromere intensities are normalized to those of RPE cells.

We have previously determined that human RPE cells have 400 molecules of CENP-A per centromere on average, equating to 200 nucleosomes in interphase (Bodor et al., 2014). By ratiometric comparison we estimate CENP-A nucleosome levels at hESCs and the two iPSC lines to be at 80, 70 and 50 nucleosomes per centromere, respectively.

We then determined the impact of the stem cell state on the larger centromere complex. Two key components of the constitutive centromere-associated network (CCAN) (Cheeseman and Desai, 2008) that make direct contacts with the kinetochore in mitosis are CENP-C and CENP-T (Gascoigne et al., 2011; Hori et al., 2008). Similar to CENP-A we find that both CENP-C and CENP-T levels are dramatically reduced at stem cell centromeres, both in embryonic-derived as well as in iPSCs (Figure 1A, B, S1C). Surprisingly, we find that the direct α-satellite binding protein CENP-B is also reduced at stem cell centromeres to 34% of RPE levels. This is unexpected as CENP-B is, at least in principle, driven by direct DNA sequence contacts (Masumoto et al., 1989).

While all centromere components analysed show reduced levels at the centromere, we find this not to be the case for the inner centromere component, Aurora B. This essential mitotic kinase (Krenn and Musacchio, 2015) is part of the chromosome passenger complex, localized to the inner centromere and important for error correction during mitosis (Carmena et al., 2012). We find Aurora B to be maintained at levels similar to somatic cells (Figures S1F and S1G), indicating that the remodelling at the centromere is unique for the kinetochore forming centromere complex.

One possible explanation for reduced centromere occupancy of CENP-A and CENP-C is that stem cells have reduced expression of centromere protein-encoding genes. To determine expression levels directly we probed extracts of RPE, hESCs, iPSCs and their parent cells for centromere protein levels. Despite reduced centromere occupancy, both embryonic and induced pluripotent stem cells maintain levels of CENP-A expression, even in excess (up to 2 fold) of those in fibroblasts, even when compared to genetically identical donor cells of iPSCs (Figure 2A, B). This is consistent with a previous report that evaluated mRNA stores of CENP-A in hESCs (Ambartsumyan et al., 2010). This uncoupling between cellular and centromeric levels in stem cells is also observed for CENP-C, where protein expression is 2 fold above that of fibroblasts. In contrast, while CENP-B is expressed in stem cells, the overall levels appear to be lower, possibly explaining the reduced centromere levels (Figure 2A, B). These results indicate that despite large pools of available CENP-A and CENP-C, these proteins are not efficiently assembled at centromeres.

**Figure 2.**
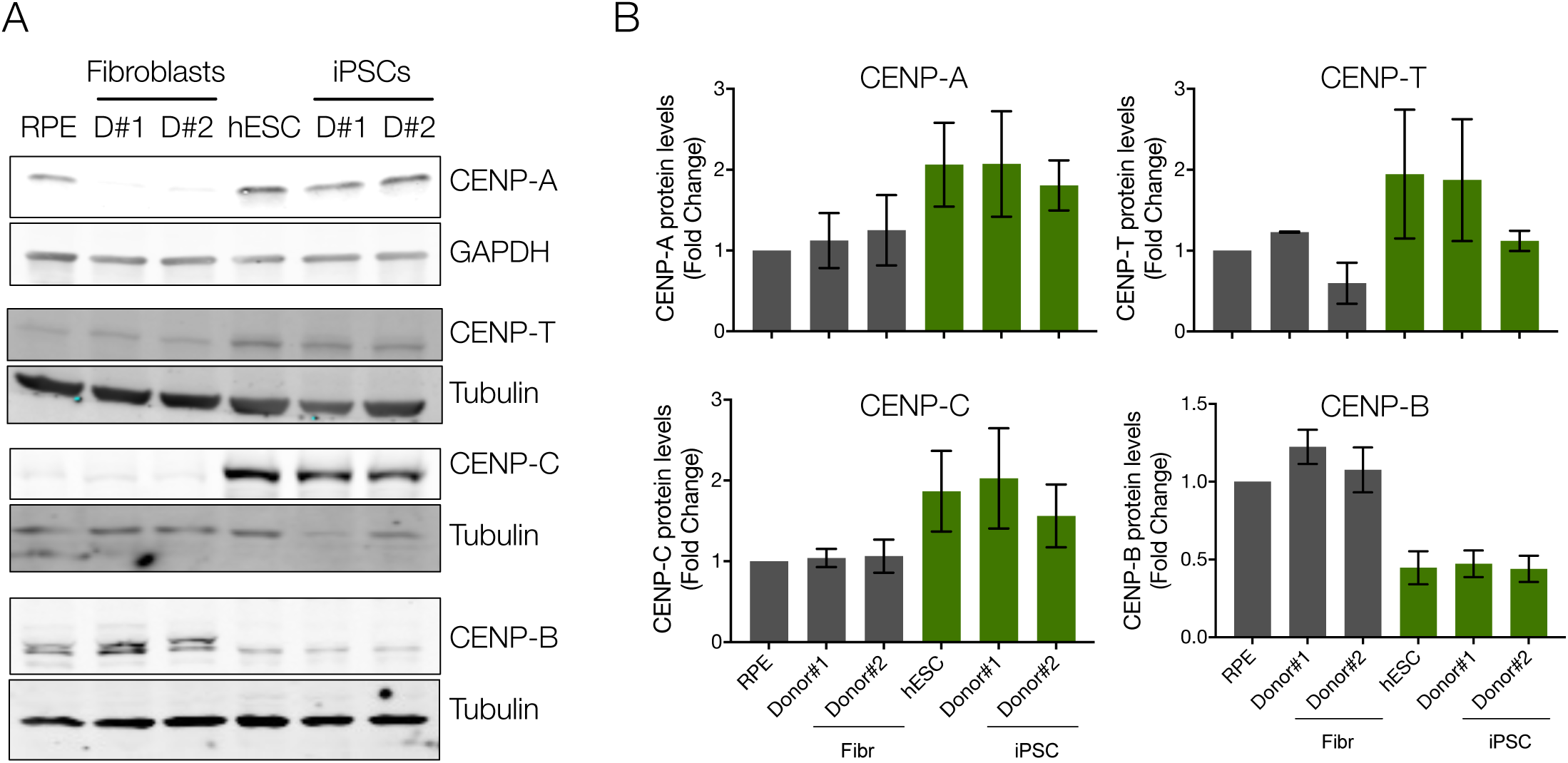
Pluripotent stem cells have elevated expression of CENP-A and CENP-C, and decreased expression of CENP-B. **A)** Human ESCs, RPE, iPSCs and the fibroblasts they were reprogrammed from, were harvested and processed for SDS-PAGE and immunobloting. CENP-A, CENP-T, CENP-C and CENP-B levels were assessed with specific antibodies. GAPDH or Tubulin were used as loading controls. **B)** Quantitation of WB bands. The average and standard error of the mean of three replicate experiments are shown. Protein levels were normalised to GAPDH or tubulin.

### CENP-A is loaded in G1 phase of the stem cell cycle

In human somatic cells, CENP-A has a unique dynamics along the cell cycle, where nucleosomes containing CENP-A are efficiently recycled on sister chromatids during S phase (Bodor et al., 2013; Jansen et al., 2007). New assembly of CENP-A occurs exclusively in early G1 phase in a CDK1 and 2 regulated manner (Jansen et al., 2007; Silva et al., 2012; Stankovic et al., 2017). Human stem cells have a characteristically abbreviated cell cycle where cells enter S phase soon after exit from mitosis (Becker et al., 2006). As G1 phase is short in these cells, CENP-A assembly dynamics could be altered. We determined the timing of CENP-A assembly using a previously established CENP-A assembly assay based on SNAP enzyme fluorescent quench-chase-pulse labelling (Bodor et al., 2012). We established a hESC line in which we introduced a SNAP-tagged CENP-A transgene by piggybac transposition to avoid gene silencing in stem cells [(Pannell et al., 2000) see methods]. We then subjected cells to a SNAP quench-chase-pulse protocol in which only nascent CENP-A-SNAP is visualised (Figure 3A). Cells were co-stained with α-tubulin to mark microtubules and identify G1 cells, based on the characteristic G1-phase-specific midbody staining. This analysis revealed that cells in G1 are positive for CENP-A assembly, similar to control somatic HeLa CENP-A-SNAP cells [Figure 3B and (Jansen et al., 2007; Silva et al., 2012; Stankovic et al., 2017)]. We therefore conclude that the G1-phase assembly is preserved in embryonic stem cells.

**Figure 3.**
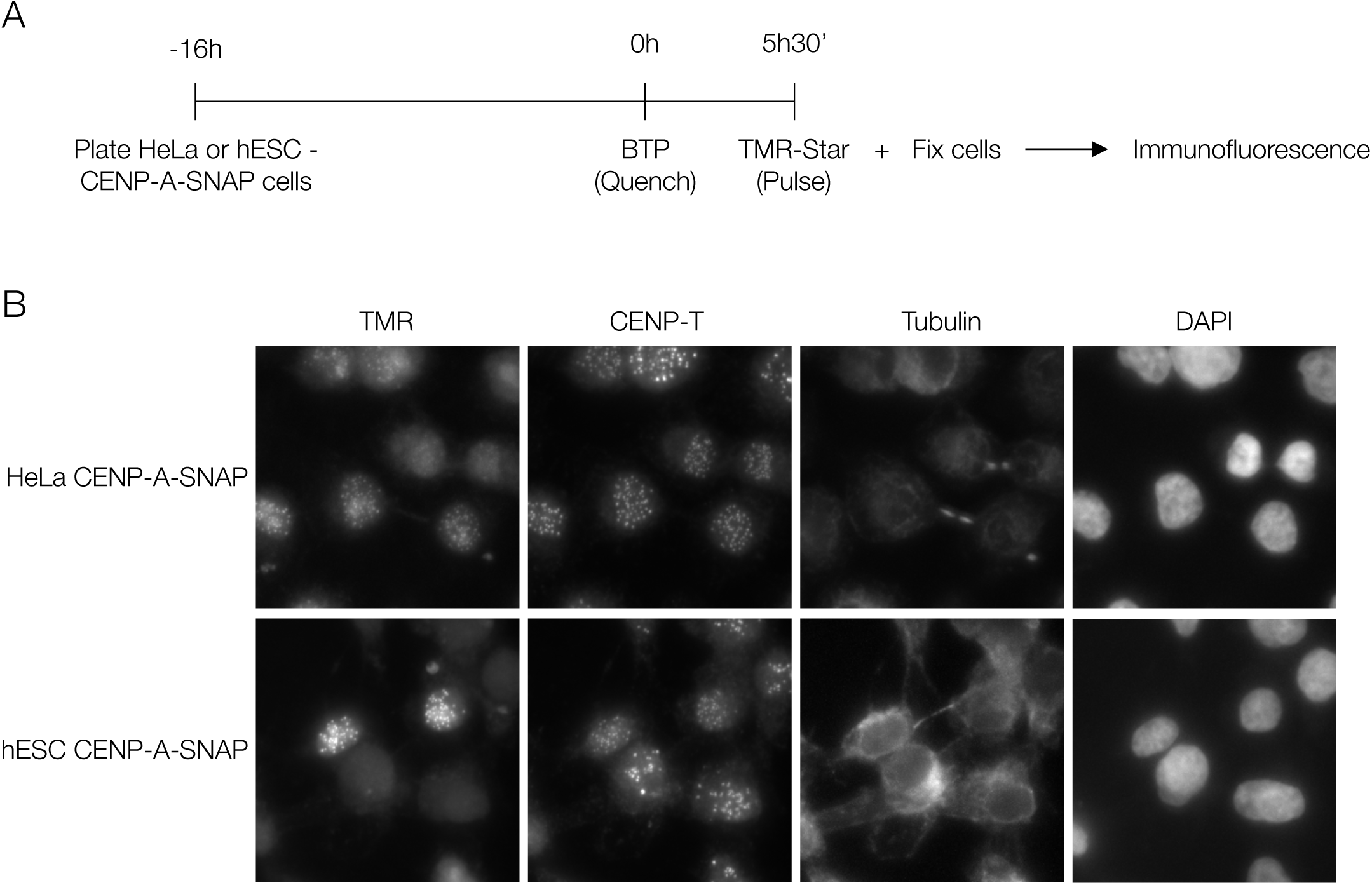
CENP-A assembles in the canonical G1 phase of the pluripotent stem cell cycle. **A)** SNAP-tag based quench-chase pulse labelling: CENP-A-SNAP expressing hESC or HeLa cells were labelled with the non-fluorescent substrate (BTP; quench) followed by a chase period (5h30min) during which new unlabelled protein is synthesised. Nascent protein is subsequently fluorescently labelled with TMR-Star (Pulse). Localization and fate of nascent fluorescently labelled CENP-A-SNAP is determined by high-resolution microscopy. **B)** Representative fluorescence images of differentiated (HeLa CENP-A-SNAP) cells or hESC (hESC CENP-A-SNAP) cells as processed according to A). Tubulin staining was used to identify midbodies, indicative of G1 phase cells. CENP-A-SNAP assembly occurs in a subset of cells and all midbody positive cells are positive for nascent CENP-A assembly.

### Mild reduction of kinetochore size of PSCs in mitosis

As we find hESCs and iPSCs to maintain a much smaller centromere complex we determined the consequences for kinetochore size which is the key protein complex to generate microtubule attachments in mitosis (Cheeseman and Desai, 2008). We stained mitotic cells for CENP-E, a mitotic kinesin, critical for chromosome congression (Wood et al., 1997). Further, we determined the levels of HEC1, an essential component of the KMN network of proteins, responsible for microtubule binding (Cheeseman et al., 2006) (Figure 4A). Both proteins accumulate on mitotic kinetochores in stem cells. While CCAN levels are low (Figure 2), both outer kinetochore components analysed are slightly reduced compared to epithelial RPE cells or donor fibroblasts (Figures 4B, C and S4A, B). Interestingly, similar to the excess cellular pools of CENP-A and CENP-C, we find that the modestly reduced kinetochore is not a consequence of a lack of expression as overall levels of both CENP-E, as well as HEC1, are higher in stem cells and iPSCs compared to RPE or fibroblasts (Figures 4D, E and S4 C, D).

**Figure 4.**
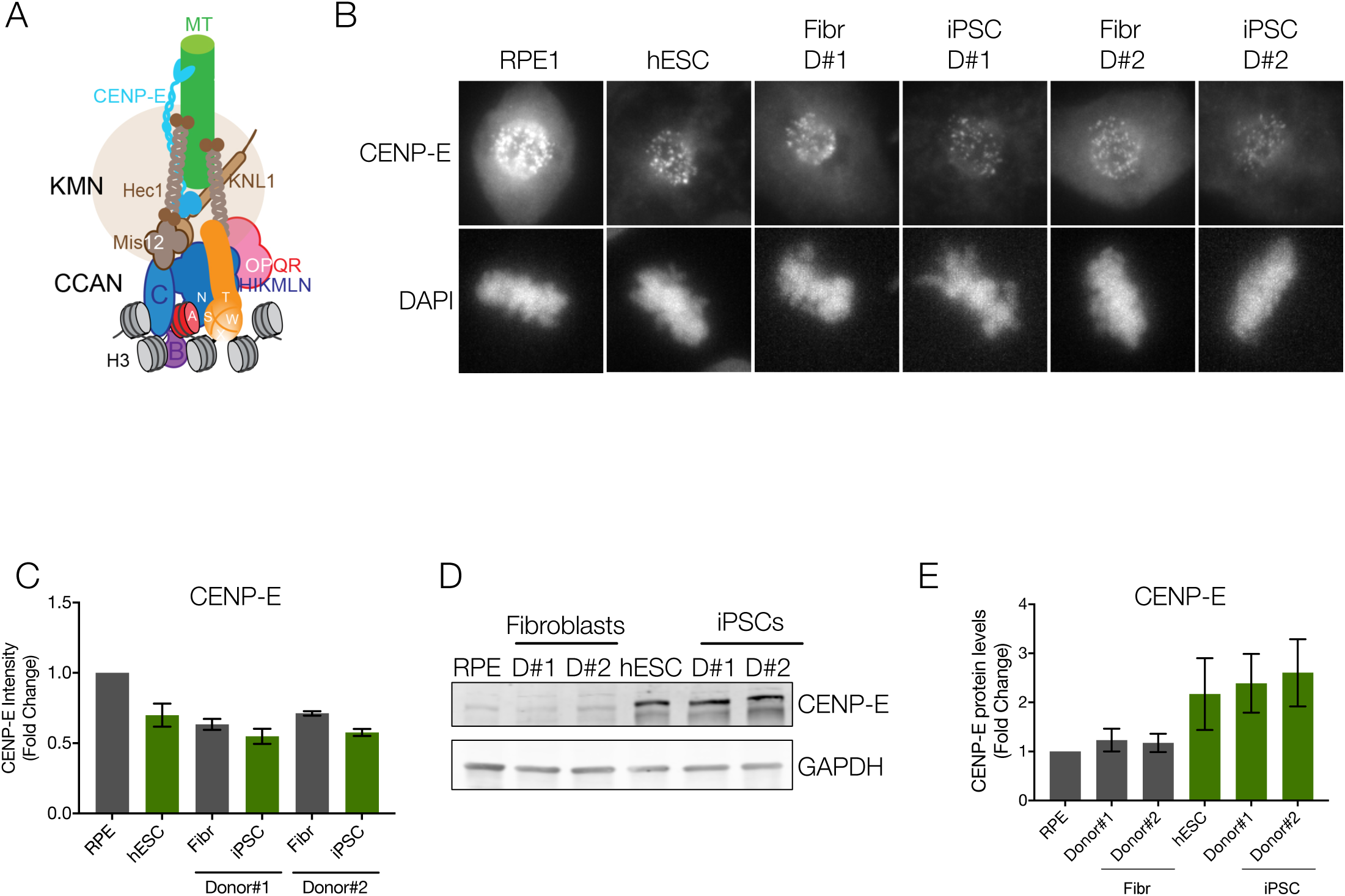
Reduced outer kinetochore size of PSCs in mitosis. **A)** Scheme representing the architecture and interactions of different proteins that comprise the human centromere and kinetochore. **B)** Representative immunofluorescence images from differentiated (RPE and Fibroblasts derived from Donor#1 and Donor#2 – Fibr D#1 and Fibr D#2) and pluripotent stem cells (human Embryonic Stem Cell line H9 - hESC and iPSCs reprogrammed from Fibr D#1 and Fibr D#2 - iPSC D#1 and iPSC D#2) for CENP-E. **B)** Quantitation of centromeric CENP-E. Mean levels of fluorescence per nuclei was measured. The average and standard error of the mean of three independent experiments are shown. **C)** Human ESCs, RPE, iPSCs and the fibroblasts they were reprogrammed from, were harvested and processed for SDS-PAGE and immunobloting. CENP-E levels were assessed with a specific antibody. GAPDH was used as a loading control. **D)** Quantitation of WB bands. Average and standard error of the mean of three independent experiments are shown. Protein levels were normalised to GAPDH.

### CENP-A loss is induced during early reprogramming of fibroblasts to iPSCs

The ability to induce the stem cell state in somatic cells offers a unique opportunity to determine the dynamics of centromeric chromatin organization and how this is linked to the formation of stem cells. The comparison of CENP-A chromatin in iPSCs and their cognate donor cells suggest that CENP-A loss is an epigenetic event that occurs during reprogramming of otherwise genetically identical cells. To determine when during the reprograming process CENP-A loss occurs we transduced fibroblasts with a cocktail of Sendai viruses expressing the four Yamanaka factors to induce pluripotency (Figure 5A). Complete iPSC formation typically requires 30 days of culturing followed by clone isolation at 40-60 days (Figure 5A). Here, we focused on very early signs of reprograming based on the expression of the pluripotency marker SSEA-4, which becomes expressed early during dedifferentiation (Chan et al., 2009). Fibroblasts do not express this cell surface protein, however they express CD13 (which is not expressed in PSCs). Taking advantage of this, we used Fluorescence-Activated Cell Sorting (FACS) to isolate SSEA-4 negative/CD13 positive (refractory to reprogramming) or SSEA-4 positive/CD13 negative (prone to reprogram) cells as early as 9 and 11 days post transduction of reprogramming factors (Figure 5A). These cells were stained for CENP-A, CENP-B and CENP-C to determine centromeric levels of the CCAN. We find that as early as 9 days, the first time point at which we can isolate a significant amount of SSEA-4 positive/CD13 negative cells, CENP-A levels show signs of decline which become more evident at 11 days post transduction (Figure 5B,C). CENP-B and to a lesser extent CENP-C levels also follow this pattern of recruitment to the centromere, with CENP-B levels decreasing as early as day 9 of reprogramming (Figure S5A, B).

**Figure 5.**
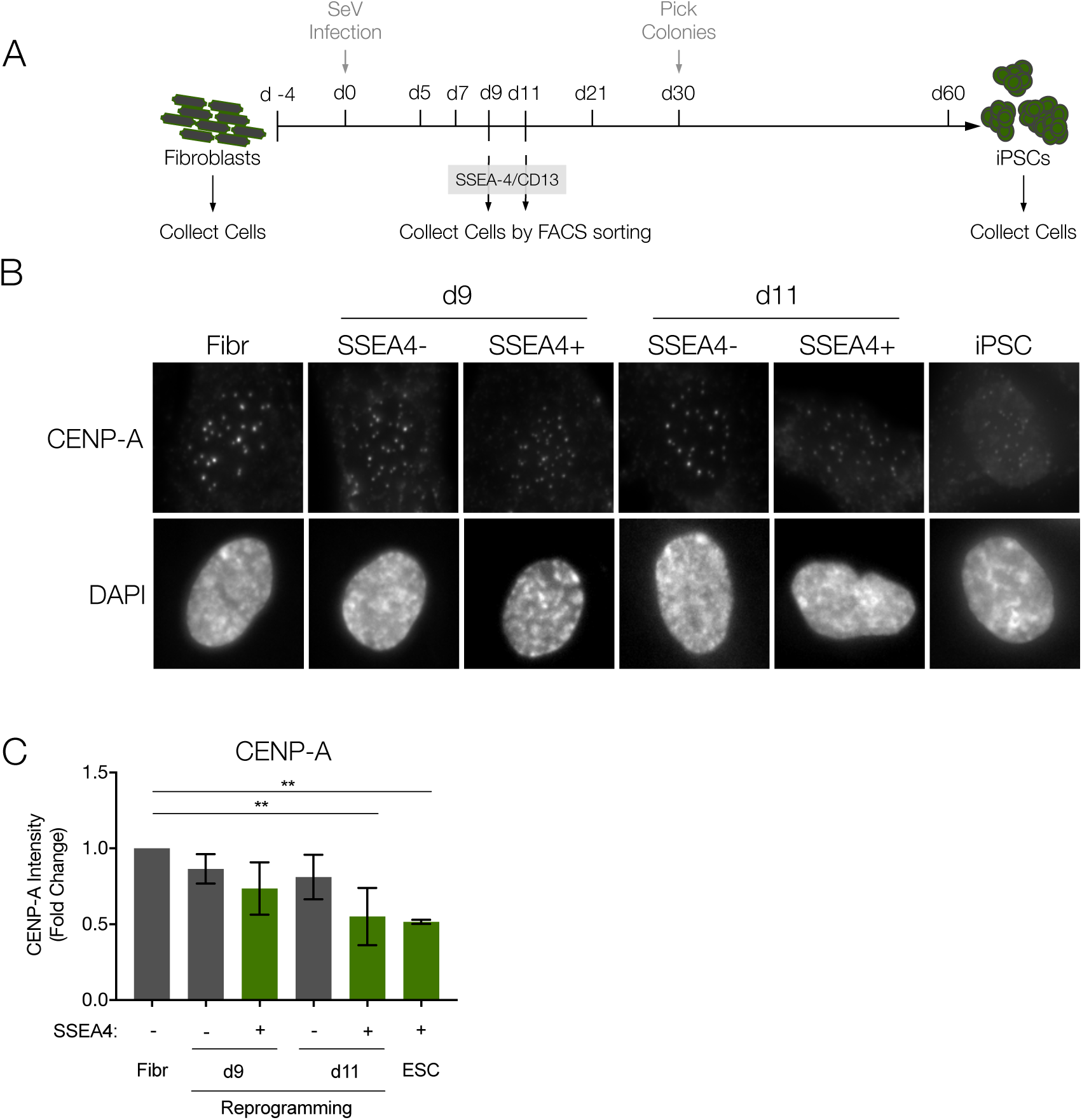
CENP-A loss is induced during early reprogramming of fibroblasts to iPSCs. **A)** Outline of the general strategy to reprogram iPSCs from fibroblasts: Human primary fibroblasts are reprogrammed by infection with Sendai Virus (SeV) expressing Oct4, Sox2, Klf4 and c-Myc. At days 9 and 11 after infection (d9 and d11, respectively), cells are incubated with antibodies specific for SSEA-4 (early pluripotency marker) and CD13 (fibroblast marker) and collected by FACS sorting. Thirty days after infection, visible colonies appear and can be picked under the microscope. Single colonies are picked, expanded and kept in culture. The cells collected at day 9 and 11, the initial fibroblast population and fully reprogrammed iPSCs (reprogrammed from those fibroblasts), were stained for CENP-A and counterstained with DAPI. **B)** Representative immunofluorescence images from cells collected by FACS at d9 and d11 and sorted by pluripotency profile (SSEA4 Negative and CD13 Positive – Refractory to reprogramming - *vs* SSEA4 positive and CD13 negative cells – Prone to reprogram) and control cells. **C)** Quantification of centromere intensities as shown in B). Average centromere intensities were determined using automatic centromere recognition and quantification (CRaQ; see methods) for indicated cell types. The average and standard error of the mean of three replicate experiments is shown for indicated cell types. Centromere intensities are normalized to those of fibroblasts.

These results indicate that the reorganization of centromeres is tightly linked to the stem cell state and correlates with early reprograming events.

## Discussion

The centromere is an essential chromosomal locus to drive chromosome segregation. While its structure and function has been studied in considerable detail in somatic, differentiated cells of different organisms, e.g. cancer cells, immortalized cells and primary cells in humans, Chicken DT40 lymphocytes and Drosophila tissue culture cells (Fukagawa and Earnshaw, 2014; McKinley and Cheeseman, 2016), relatively little is known about centromere structure in stem cell populations. Aspects of centromere biology have been reported in stem cells of the Arabidopsis meristem and Drosophila midgut and male germline (García Del Arco et al., 2018; Lermontova et al., 2006; Ranjan et al., 2018) but centromere structure and size has not been thoroughly investigated in those systems.

Using human embryonic stem cells and iPSCs as a model we found that these cells maintain a low level of centromeric chromatin as well as associated centromere proteins, despite abundant cellular pools. Interestingly, the inner centromere component Aurora B is maintained at normal levels and does not seem affected in PSCs. Moreover, we find that the weak centromere seems to only moderately affect the recruitment of kinetochore proteins in mitosis. These findings indicate that CCAN size and kinetochore size regulation can be uncoupled, and that stem cells have the ability to partially, but not fully, compensate for the reduced centromeric chromatin size. Although this does not seem to be a conserved characteristic of the centromere (Drpic et al., 2018), we previously showed this to be the case in RPE cells in which forced reduction or expansion of CENP-A chromatin had little impact on kinetochore size (Bodor et al., 2014). We now find a physiological example of a partial compensatory mechanism within the kinetochore.

It has previously been shown that, in Drosophila, CENP-A assembles in telophase/early G1 in brain stem cells (Dunleavy et al., 2012). An increase in CENP-A in G2 in germline stem cells has also been suggested recently (Ranjan et al., 2019). Here we show that assembly of CENP-A chromatin occurs in G1 phase of the stem cell cycle, as is the case in human differentiated and immortalized cells and in cancer cell lines (Jansen et al., 2007; Silva et al., 2012). An open question remains how CENP-A levels are restricted in stem cells. One possibility is that cells that exit mitosis and rapidly transition into S phase have a relatively short G1 phase window during which CENP-A can be assembled before inhibitory Cdk activity rises (Silva et al., 2012). It is tempting to speculate that this combined with the lack of CENP-B could lead to the destabilisation of CENP-A and CENP-C (Fachinetti et al., 2015), resulting in a weaker centromere in PSCs.

We further find that reduction in centromeric chromatin size is induced early during iPSC reprogramming, coincident with the time of cell cycle shorting. Profound remodelling of chromatin marks is observed during reprogramming and one of the earliest events in reprogramming is the rapid genome-wide re-distribution of H3K4me2 during both mouse and human somatic cell reprogramming (Cacchiarelli et al., 2015; Koche et al., 2011). Moreover, methylation of H3K4me2 by Wrd5 to a trimethylated state, leading to a global decrease in di-methylation, is required for both self-renewal and efficient reprogramming of somatic cells (Ang et al., 2011). H3K4me2 depletion at engineered centromeric chromatin causes defects in HJURP recruitment and CENP-A assembly and consequent kinetochore dysfunction and chromosome missegregation (Bergmann et al., 2011). These and other major chromatin changes that also occur during this early window, including DNA methylation erasure, could play a role in CENP-A chromatin remodelling.

Finally, cultured stem cells are prone to chromosome missegregation compared to somatic cells. While this can be a consequence, at least in part, of cell culture conditions, our findings that stem cells maintain a reduced centromere complex, may impact on chromosome segregation fidelity. It will be interesting to establish whether there is a causal link in stem cell centromere functionality and the tendency of these cells to missegregate.

## Methods

### Cell culture

All cell lines were grown at 37°C in 5% CO2 incubators. Normal human dermal fibroblasts (NHDF - GIBCO) were maintained in fibroblast medium (DMEM high glucose, 10% FBS, 1% Pen-Strep, 1% MEM Non-Essential Amino Acids and 50 μM 2-mercaptoethanol). H9 ESC (hESC) and hiPSC lines were grown in VTN coated plates in Essential-8 medium (TeSR-E8, Stem Cell Technologies), and dissociated with gentle cell dissociation reagent (0.5mM EDTA in PBS) or Tryple-Express Enzyme (Gibco) when single cell dissociation was necessary. RPE-1 cells were grown in RPE medium (DMEM/F-12, 10%FBS, 1% Pen-Strep, 2mM L-Glutamine, 1.6% Sodium bicarbonate). HeLa-CENP-A SNAP clone #72 (Jansen et al., 2007) was grown in HeLa medium (DMEM high glucose, 10% FBS, 1% Pen-Strep, 2mM L-Glutamine).

### Reprogramming of human Fibroblasts to iPSCs

Reprogramming was performed as described previously (Milagre et al., 2017). Briefly, 3.0×10^5^ NHDFs were transduced with CytoTune®-iPS 2.0 Sendai Reprogramming Kit (Invitrogen), according to manufacturer’s instruction, at an MOI of 1. Cells were maintained in fibroblast medium (DMEM, 10% FBS, 1% Pen-Strep, 1% MEM Non-Essential Amino Acids and 50 µM 2-mercaptoethanol) for five days. Transduced cells were then replated onto VTN (Invitrogen) coated dishes and maintained in Essential 8 medium (E8 - stem cell technologies). Medium was replenished daily. Cells were collected at different time-points during reprogramming by FACS (d9, d11) or manually (NHDFs and fully established iPSCs).

### Immunofluorescence, microscopy and image analysis

Cells were grown on glass coverslips coated with poly-L lysine (Sigma-Aldrich) or VTN (Thermo Fischer Scientific) and fixed with 4% formaldehyde (Thermo Scientific) for 10 min followed by quenching with 100mM Tris-HCl. Cells were permeabilised in PBS with 0.3% Triton-X-100. All primary antibody incubations were performed at 37°C for 1h in a humid chamber. Fluorescent secondary antibodies were from Jackson ImmunoResearch (West Grove, PA) or Rockland ImmunoChemicals (Limerick, PA) and used at a dilution of 1:250. All secondary antibody incubations were performed at 37°C for 45 min in a humid chamber. Cells were counter-stained with DAPI (4’,6-diamidino-2-phenylindole; Sigma-Aldrich) before mounting in Mowiol.

The following primary antibodies and dilutions were used: mouse monoclonal anti-CENPA (#ab13939, abcam) at 1:500, rabbit polyclonal anti-CENP-B (#ab25734, Abcam) at 1:500, guinea-pig polyclonal anti-CENP-C (#PD030, MBL International) at 1:1000, rabbit polyclonal anti-CENP-T at 1:250 (#ab220280, Abcam), goat anti-Sox2 (#AF2018, R&D) at 1:200, goat anti-Nanog (#AF1997, R&D) at 1:100, rabbit anti-CENP-E (kind gift from Don Cleveland) at 1:200, mouse monoclonal anti-Aurora B (#611082, BD Transduction Laboratories) at 1:200, mouse monoclonal anti-HEC1 (Thermo Scientific Pierce MA1-23308) at 1:100 and rat monoclonal anti-Tubulin (SC-53029, Santa Cruz Biotechnology, Dallas, TX) at 1:10,000.

Z-stack slices were captured with wide field microscopes, either Leica High Content Screening microscope, based on Leica DMI6000 equipped with a Hamamatsu Flash Orca 4.0 sCMOS camera, using a 63x oil objective (HC PLAN APO, NA 1.4) with 0.2 µm z sections, or Deltavision Core system (Applied Precision) inverted microscope (Olympus, IX-71) coupled to Cascade2 EMCCD camera (Photometrics), using a 60x oil objective (Plan Apo N, NA 1.42) with 0.2 µm z sections.

Immunofluorescent signals were quantified using the CRaQ (Centromere Recognition and Quantification) method (Bodor et al., 2012) using CENP-A, CENP-T or CENP-C as centromeric reference. Alternatively, Hec1 and CENP-E levels were measured only in mitotic cells using an ImageJ based macro, which measures the median intensity of the whole nucleus.

### Western blot (WB) analysis

For WB analysis, whole cell extracts were resolved by SDS-PAGE and blotted onto Nitrocellulose membranes. Membranes were blocked in TBS-Tween (10% powdered milk) or Odyssey blocking buffer (Li-cor Biosciences) and incubated overnight at 4°C with the indicated antibodies. Secondary antibodies were used at 1:10000 prior to detection on Odyssey near-infrared scanner (Li-cor Biosciences).

The following primary antibodies were used for WB: rabbit polyclonal anti-CENP-A (#2186, Cell Signaling Technology) at 1:500, rabbit polyclonal anti-CENP-B (#ab25734, Abcam) at 1:200,rabbit polyclonal anti-CENP-T (#ab220280, Abcam) at 1:250, guinea-pig polyclonal anti-CENP-C (#PD030, MBL International) at 1:250, rabbit polyclonal anti-H4K20me (#ab9052, Abcam) at 1:4000, rabbit anti-CENP-E (kind gift from Don Cleveland) 1:250, mouse monoclonal anti-Hec1 (#MA1-23308, Thermo Fischer Scientific) at 1:250, mouse monoclonal anti-α tubulin (T9026, Sigma-Aldrich) at 1:5000, rabbit monoclonal anti-GAPDH (#2118S, Cell Signaling) at 1:2000. Secondary antibodies used: IRDye800CW anti-rabbit (Li-cor Biosciences), IRDyLight800CW anti-rabbit (Li-cor Biosciences), IRDyLight800CW anti-guinea (Li-cor Biosciences), IRDyLight800CW anti-mouse (Li-cor Biosciences) and IRDyLight680LT anti-mouse (Li-cor Biosciences).

### Cell Fractionation

Cell fractionation was performed for RPE and hESC lines after cell lysis in ice cold buffer [50 mM Tris-HCl (pH 7.5), 150 mM NaCl, 0.5 mM EDTA, 1% Triton-X 100, and a protease inhibitor cocktail (ROCHE)]. Soluble proteins were separated from the insoluble fraction by centrifugation at 21,000×g at 4°C and resuspended in an equal volumes of lysis buffer. Pellet fraction was incubated with 1.25 U/μl of benzonase nuclease (Merck, Millipore, Burlington, MA) on ice for 10 min prior to denaturation in 4X loading buffer (Li-Cor).

### DNA constructs

To obtain the hESC CENP-A-SNAP cell line we re-cloned CENP-A-SNAP, from pBABE-CENP-A SNAP plasmid (Jansen et al., 2007), to avoid retroviral silencing, onto a piggybac plasmid (pB-CAG-Dest-pA-pgk-bsd - kind gift from José Silva).

### Stable cell lines

hESC H9 cell line was transfected with 2ug of pB-CAG-Dest-pA-pgk-bsd-CENP-A-SNAP plus 2ug of pBASE plasmid (harbouring the piggybac transposase, kind gift from José Silva) using FuGeneHD (Roche), in a ratio of DNA:FuGene of 1:3. Cells were then subjected to 5 days blasticidin selection and single clones were picked and characterised for CENP-A-SNAP protein levels by western blot.

### Quench-chase-pulse labelling

Cell lines expressing CENP-A-SNAP were quench-pulse labelled as previously described (Bodor et al., 2012). Briefly, cells were quenched with a non-fluorescent bromothenylpteridine (BTP; New England Biolabs) at 2 µM final concentration, and kept in culture for 5 hours and 30 minutes. Cells were then pulse labelled with tetra-methyl-rhodamine-conjugated SNAP substrate (TMR-Star; New England Biolabs) at 4 µM final concentration, labelling all newly synthesised CENP-A molecules at the centromere, and fixed for immunofluorescence.

### Fluorescence-activated cell sorting (FACS)

For cell sorting, cells undergoing reprogramming were incubated with antibodies against CD13 (PE, BD Pharmigen) and SSEA-4 (Alexa Fluor 647, BD Pharmigen) for 30 min. Cells were washed in a 2% FBS/PBS solution and passed through a 50µm cell strainer to obtain a single-cell suspension.

Appropriate negative and positive controls were used to assess optimal FACS conditions. Cell sorting was performed using a FACSAria cell sorter instrument (BD Biosciences) and cells were collected for immunofluorescence.

## Author Contributions

I.M. conceived the project, designed and performed experiments, analysed data, and wrote the manuscript; C.P. performed western blot and cell fractionation experiments, R.A.O and L.J. interpreted the data, provided helpful discussions for project design and wrote the manuscript. All authors have interpreted the data and provided helpful discussions, read and approved the manuscript.

## Acknowledgements

We thank Don W. Cleveland (UCSD) for CENP-E antibodies, Jose Silva for the pB-CAG-Dest-pA-pgk-bsd and pBASE plasmids and João Mata (IGC) for technical support. We would like to thank the members of the Jansen and Oliveira Labs for helpful discussions. We would like to thank the technical support of IGC’s Advanced Imaging Facility (AIF-UIC), which is supported by the national Portuguese funding ref# PPBI-POCI-01-0145-FEDER-022122, co-financed by Lisboa Regional Operational Programme (Lisboa 2020), under the Portugal 2020 Partnership Agreement, through the European Regional Development Fund (FEDER) and Fundação para a Ciência e a Tecnologia (FCT; Portugal). We would like to thank the Flow Cytometry Facility of Instituto Gulbenkian de Ciência for their services and assistance. This work was funded by Fundação para a Ciência e Tecnologia project: PTDC/BIA-CEL/31502/2017 and was supported by the European Union’s Horizon 2020 research and innovation programme under the Marie Skłodowska-Curie grant agreement No 704763 to (Marie Sklodowska-Curie fellowship to IM) as well as funding from an ERC-consolidator grant ERC-2013-CoG-615638 and a Wellcome Senior Research Fellowship in Basic Biomedical Science to LETJ and EMBO Installation Grant IG2778 to RAO.

---

**Figure S1.**
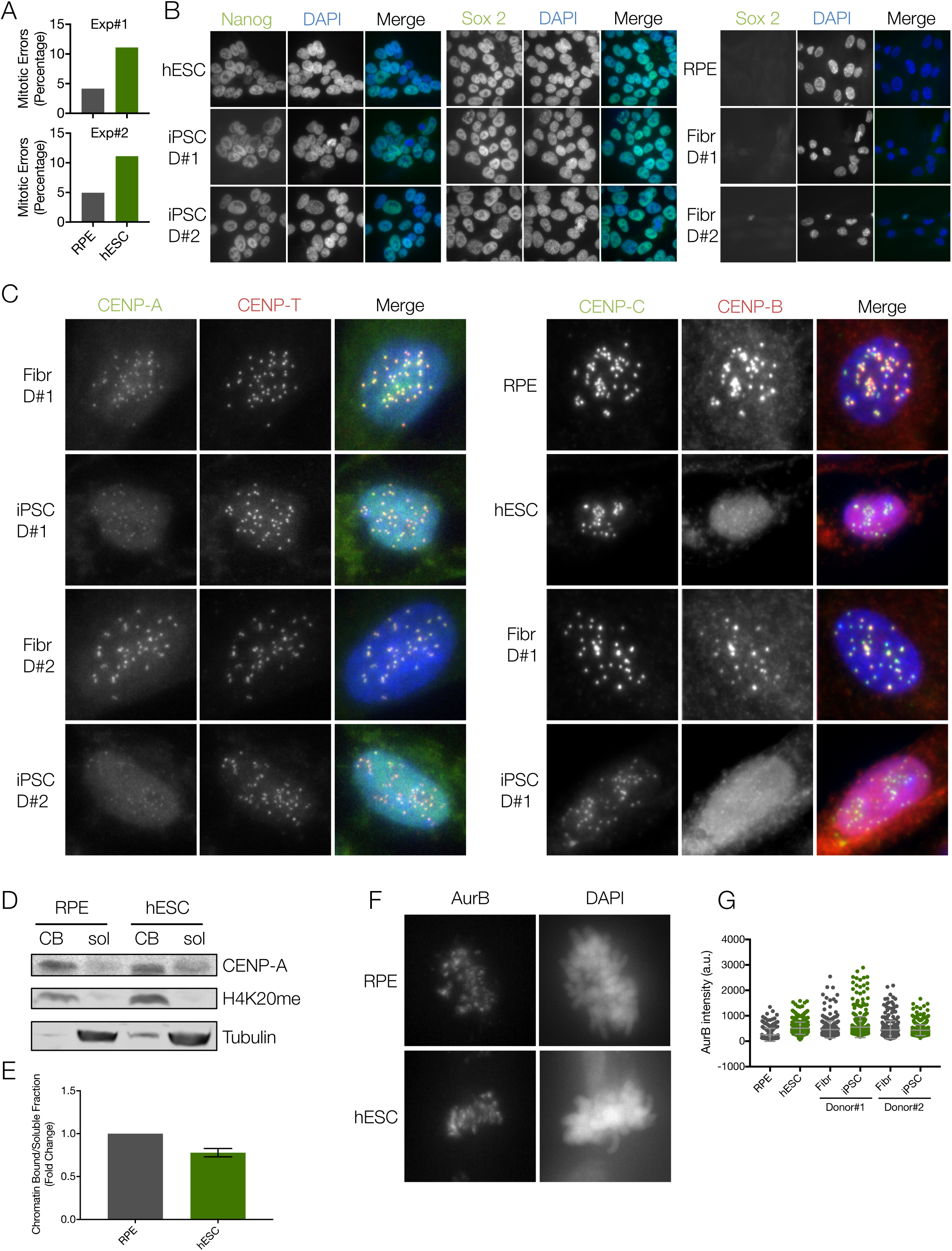
Pluripotent stem cells have increased errors, a weaker centromere, but normal levels of AurB, when compared to differentiated cells. **A)** Quantification of mitotic errors in RPE and hESC, from two independent experiments. Cells were fixed and the frequency of mitotic errors in unperturbed cells was evaluated. **B), C)** Differentiated (RPE and Fibr D#1 and Fibr D#2) and pluripotent stem cells (hESC or iPSC D#1 or iPSC D#2) were fixed and stained for either B) Nanog or Sox2 and counterstained with DAPI. Representative immunofluorescence images are shown, or C) CENP-A, CENP-T, CENP-C or CENP-B and counterstained with DAPI. Representative immunofluorescence images from Fibroblasts and iPSC from Donor #1 and Donor #2 are shown for CENP-A and CENP-T and representative images from RPE and hESC, Fibroblasts and iPSC from donor #1 are shown for CENP-B and CENP-C. **D)** and **E)** Cell fractionation experiments to assess total levels of soluble and chromatin bound CENP-A in RPE and hESC. Immunoblot probed for soluble (sol) and chromatin bound (CB) fractions of CENP-A in RPE and hESC. Tubulin is used as a marker for the soluble fraction and histone H4K20me2 for the CB fraction (D). Quantification of CENP-A ratio (chromatin bound/soluble fraction) from three independent experiments (E). **F)** Differentiated (RPE and Fibr D#1 and Fibr D#2) and pluripotent stem cells (hESC or iPSC D#1 or iPSC D#2) were fixed and stained for Aurora B (AurB) and counterstained with DAPI. **G)** Quantification of centromere intensities for AurB. Average centromere intensities were determined using automatic centromere recognition and quantification (CRaQ) for indicated cell types. Horizontal lines represent the mean for each sample.

**Figure S4.**
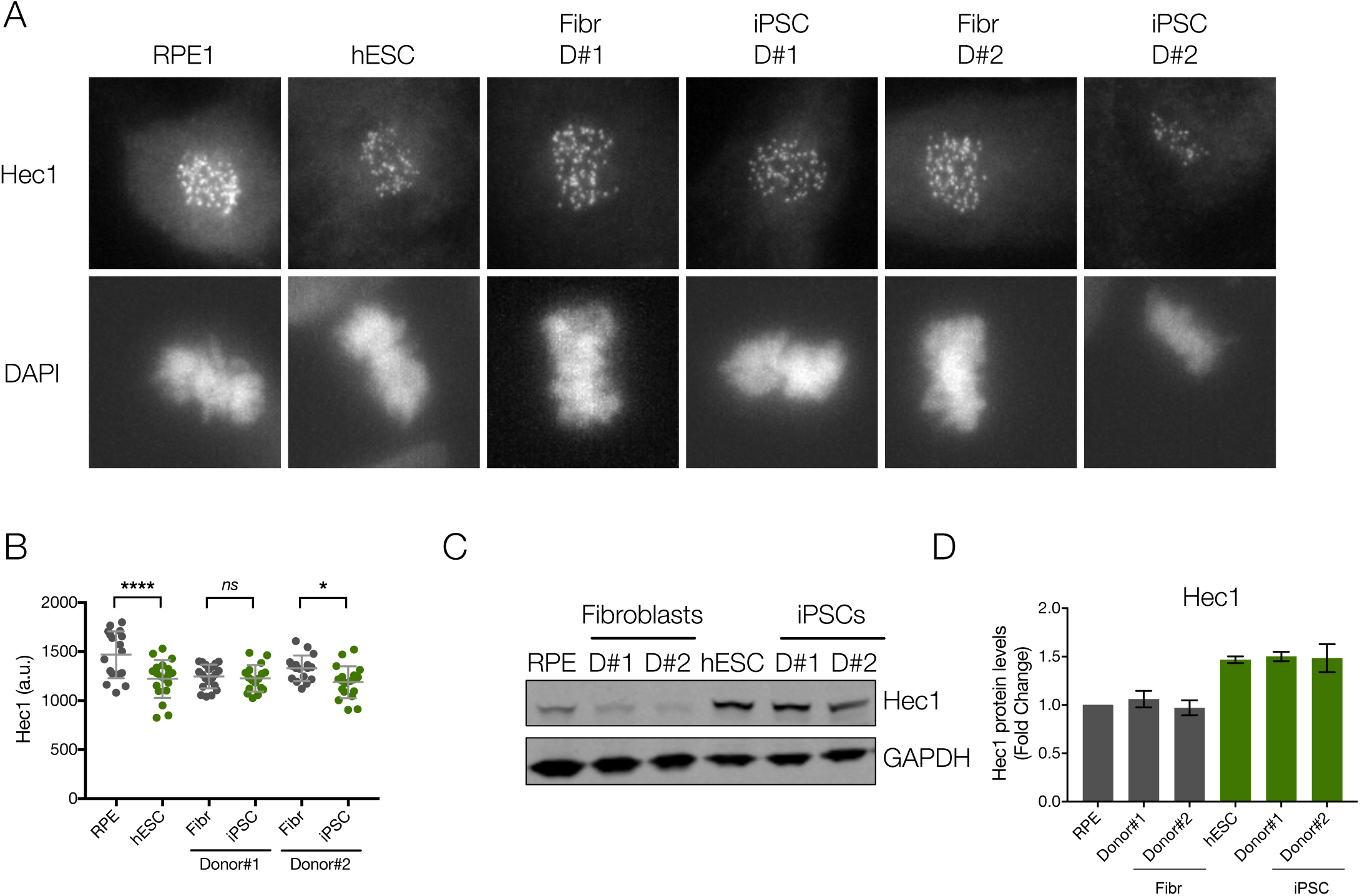
Reduced Hec1 at the kinetochore of PSCs in mitosis. **A)** Representative immunofluorescence images from differentiated (RPE and Fibroblasts derived from Donor#1 and Donor#2 – Fibr D#1 and Fibr D#2) and pluripotent stem cells (human Embryonic Stem Cell line H9 - hESC and iPSCs reprogrammed from Fibr D#1 and Fibr D#2 - iPSC D#1 and iPSC D#2) for Hec1. **B)** Quantitation of centromeric Hec1. Mean levels of fluorescence per nuclei was measured. Horizontal lines represent the mean, whiskers represent standard deviation, for each sample. **C)** Human ESCs, RPE, iPSCs and the fibroblasts they were reprogrammed from, were harvested and processed for SDS-PAGE and blotted for Hec1 levels. GAPDH was used as a loading control. **D)** Quantitation of WB bands. Average and standard error of the mean of three independent experiments are plotted. Protein levels were normalised to GAPDH.

**Figure S5.**
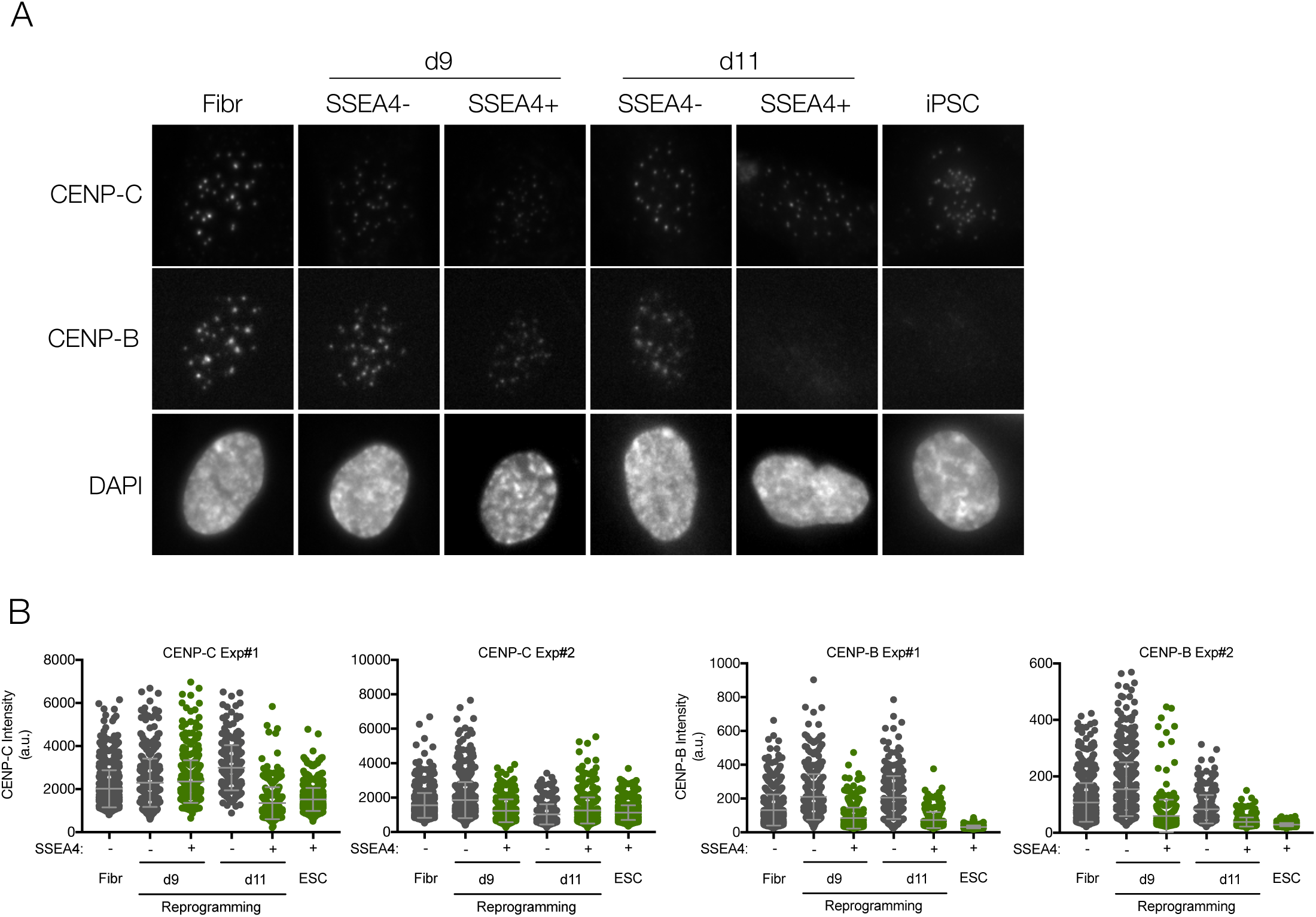
CENP-B and CENP-C are reduced during early reprogramming of fibroblasts to iPSCs. **A)** Experiment as in Figure 5. Representative immunofluorescence images from cells collected by FACS at d9 and d11 and sorted by pluripotency profile as in Figure 5, stained with CENP-C and CENP-B and counterstained with DAPI. **B)** Quantification of centromere intensities as shown in A). Average centromere intensities were determined using automatic centromere recognition and quantification (CRaQ) for indicated cell types. Horizontal lines represent the mean, whiskers represent standard deviation, for each sample.

